# Peptide-based epitope design on non-structural proteins of SARS-CoV-2

**DOI:** 10.1101/2021.12.27.474315

**Authors:** R S Swathika, S Vimal, E Bhagyashree, Elakkiya Elumalai, Krishna Kant Gupta

## Abstract

The SARS-CoV-2 virus has caused the severe pandemic, COVID19 and since then its been critical to produce a potent vaccine to prevent the quick transmission and also to avoid alarming deaths. Among all type of vaccines peptide based epitope design tend to outshine with respect to low cost production and more efficacy. Therefore, we started with obtaining the necessary protein sequences from NCBI database of SARS-CoV-2 virus and filtered with respect to antigenicity, virulency, pathogenicity and non-homologous nature with human proteome using different available online tools and servers. The promising proteins was checked for containing common B and T-cell epitopes. The structure for these proteins were modeled from I-TASSER server followed by its refinement and validation. The predicted common epitopes were mapped on modeled structures of proteins by using Pepitope server. The surface exposed epitopes were docked with the most common allele DRB1*0101 using the GalaxyPepDock server. The epitopes, **ELEGIQYGRS** from Leader protein (NSP1), **YGPFVDRQTA** from 3c-like proteinase (nsp5), **DLKWARFPKS** from NSP9 and **YQDVNCTEVP** from Surface glycoprotein (spike protein) are the epitopes which has more hydrogen bonds. Hence these four epitopes could be considered as a more promising epitopes and these epitopes can be used for future studies.

## Introduction

In December 2019, COVID-19 was first reported in Wuhan, China. It is a severe acute respiratory syndrome that causes respiratory disease which leads to death or other serious complications such as heart disorders and multi organ failure [1]. The virus was first transmitted from animals to humans. Majority of infected people shows moderate symptoms but pathological effects of COVID-19 is heterogeneous. Person to person transmission occurs through contact with infected secretion mainly via respiratory droplets.

Currently, several old drugs such as chloroquine (antimalarial), remdesivir(antiviral), azithromycin (antibiotic), tocilizumab (immunomodulatory agent) and corticosteroids, are being used to counteract the symptoms of COVID-19 [2,3,4]. The traditional vaccines have to many disadvantages such as allergic and autoimmune reactions. They need storage at a cold temperature and have low stability and. Therefore, peptide-based vaccines may be a promising strategy which involves minimal microbial components to stimulate humoral and adaptive immunity against a microorganism. These vaccines are able to target very specific epitopes removing the risks associated with allergic and autoimmune responses [5].

The SARS CoV 2 contains four major structural proteins: Membrane protein, Spike protein, Envelope protein and Nucleo-capsid protein. Early work on the new corona virus have highlighted the importance of surface glycoprotein (spike protein), also called as S proteins which act as the key mediator for the virus to enter the host cell [6]. While comparing SARS COVID and SARS COVID2 the spike protein is found to be binding to a receptor called angiotensin converting receptor ACE-2 which helps in entering to the host cells. It has been reported that the spike protein of the novel virus binds to ACE -2 more efficiently when compared to that of SARS COVID. Therefore, it spreads more effortlessly and is also more contagious. Many major pharmaceutical industries and also universities with research facilities are working hard to find a potential vaccine to curb the global pandemic situation [7,8].

Peptides have become one of the most important vaccine candidates given their comparatively easy production and design, chemical stability of structure and non-appearance of any possible infectious potential[9,10,11,12]. Hence we used substractive genomics and reverse vaccinology [13] approach to contribute towards peptide based epitope design to curb this pandemic infection. All 38 proteins (Genbank accession number: NC_045512) encoded by SARS CoV 2 genome was screened for its Antigenicity, essentiality, virulency and non-homology with human proteome. The final set of proteins were screened for B and T cell epitopes, mapped on 3D protein structures and finally docked with DRB10101. Parallel bionformatic predictions identified a priori potential B and T cell epitopes for 2019-nCoV. We hope that there is a high probability that these epitopes are targets for immune recognition of 2019-nCoV.

## 2. METHODOLOGY

### 2.1. Retrieval of proteome sequence data

The protein sequence of the SARS-CoV-19 NC_045512 was retrieved from NCBI (Genbank accession number: **NC_045512**). It consists of the complete proteome sequence of SARSCoV-19. The FASTA format for all proteins were retrieved.

### 2.2. Comparative analysis with human genome

Each protein sequence was compared with human proteome by using **BLASTp** to check the non-homogeneity to avoid auto immunity. BLASTP programs search protein databases using a protein query.

### 2.3. Antigenicity and virulence prediction

The non-homologous proteins must be antigenic for the stimulation of both B and T cell immunity. Therefore, the antigenicity and virulency were predicted for those proteins using the **Vaxijen2.0** [14] and **MP3** [15] servers respectively.

### 2.4. Position of protein sequences in the transmembrane region

The **TMHMM** server [16] was used to analyse the trans-membrane position of protein. It is a membrane protein topology prediction method based on a hidden Markov model.

### 2.5. Evaluation of essential proteins

The proteins were checked for its essentiality to the virus by using database of essential gene (**DEG ver.15.2**). DEG [17] is a database that consists of data which includes protein coding genes, on-coding RNA and other essential genomic elements. **BLASTp** with default parameters were used to find essential proteins.

### 2.6. Epitope prediction

B and T cells are the most responsible for inducing immune response in human body. For the obtained protein sequences B cell epitopes were found using **ABCpred server** [18]. To determine each sequences’ binding affinity towards MHC-I and MHC-II molecules, **MHC-I binding prediction tool** (**http://tools.immuneepitope.org/mhci/**) and **MHC-II binding prediction tool** (**http://tools.iedb.org/mhcii/**) were used respectively. A quantitative prediction server MHCpred v.2.0 [19] was used to find half maximal (50%) inhibitory concentration IC_50_. The epitopes which had IC_50_ values comparatively less than 100 nM were taken for further process.

### 2.7. Prediction and analysis of 3D structure

The 3D structure of the target proteins was predicted using online software I-TASSER [20]. It builds three dimensional atomic models from multiple threading alignments and iterative structural assembly simulations. The obtained structures were refined using GALAXYWEB server [21] and the appropriate models were made choice by considering highest C-score value. The selected models were validated using the SAVES server (**https://saves.mbi.ucla.edu/**) which includes PROCHECK which estimates the dependability of the proteins by calculating the proteins’ stereo chemical quality which is done by constructing the Ramachandran plot for the backbone structure of the protein.

### 2.8. Epitope exo-membrane topology evaluation

The exo-membrane topology of the proteins were analysed using the Pepitope server (**http://pepitope.tau.ac.il/**) where position of each epitope in the predicted protein structure model was identified. Under epitope mapping algorithm, “**Combined**” option was selected. This is done to make sure that the targeted epitopes are exposed on the surface and ensure that the epitopes considered are not hidden within the protein globular structure.

### 2.9. Epitope binding mode and interaction analysis

The selected epitopes which fulfilled all the constrains were then subjected to docking with the 3-D structure of most dominant allele DRB1*0101 (1AQD). To perform docking **GALAXY Pepdock server** [22] was used. From the obtained peptide-protein complex, the best complex was considered based on hydrogen bonding pattern, similarity score and accuracy score. The binding mode of the complex and binding interaction was visualised using the **LIGPLOT tool** [23].

## 3. RESULTS AND DISCUSSION

### 3.1 Sequence retrieval and evaluation of protein sequences

The Primary genomic sequence data of SARS-CoV-19 was fetched from the NCBI database. From the obtained Genomic data 38 protein sequences were collected in FASTA format.

### 3.2 Virulency

Virulency is considered as one of the main factor for studying the pathogenesis and the MP3 tool uses a combined SVM-HMM approach to yield increased efficiency and reliability to predict pathogenic proteins. Out of 38 proteins, 11 proteins were found to be virulent (Table 1 & Supplementary Table1).

**Table1:** Final set of nCoV proteins for epitope prediction.

| **Locus** | **Protein name** | **Human protein Similarity** | **Virulence** | **Antigenicity** | **TM Helix/Outside region** | **Essentiality of protein** |
| --- | --- | --- | --- | --- | --- | --- |
| **YP_009725297** | Leader protein(nsp1) | No | P | 0.4064 | Outside | Yes |
| **YP_009725301** | 3c-like proteinase(nsp5) | No | P | 0.4159 | Outside | Yes |
| **YP_009725304** | nsp8 | No | P | 0.4008 | Outside | Yes |
| **YP_009725305** | nsp9 | No | P | 0.6476 | Outside | Yes |
| **YP_009725306** | nsp10 | No | P | 0.4039 | Outside | Yes |
| **YP_009742608** | Leader protein | No | P | 0.4064 | Outside | Yes |
| **YP_009742612** | 3c-like proteinase | No | P | 0.4078 | Outside | Yes |
| **YP_009742615** | nsp8 | No | P | 0.4107 | Outside | Yes |
| **YP_009742616** | nsp9 | No | P | 0.6226 | Outside | Yes |
| **YP_009742617** | nsp10 | No | P | 0.4149 | Outside | Yes |
| **YP_009724390** | Surface glycoprotein(spike protein) | No | P | 0.4661 | Outside/TM helix | Yes |

**Table 2:**
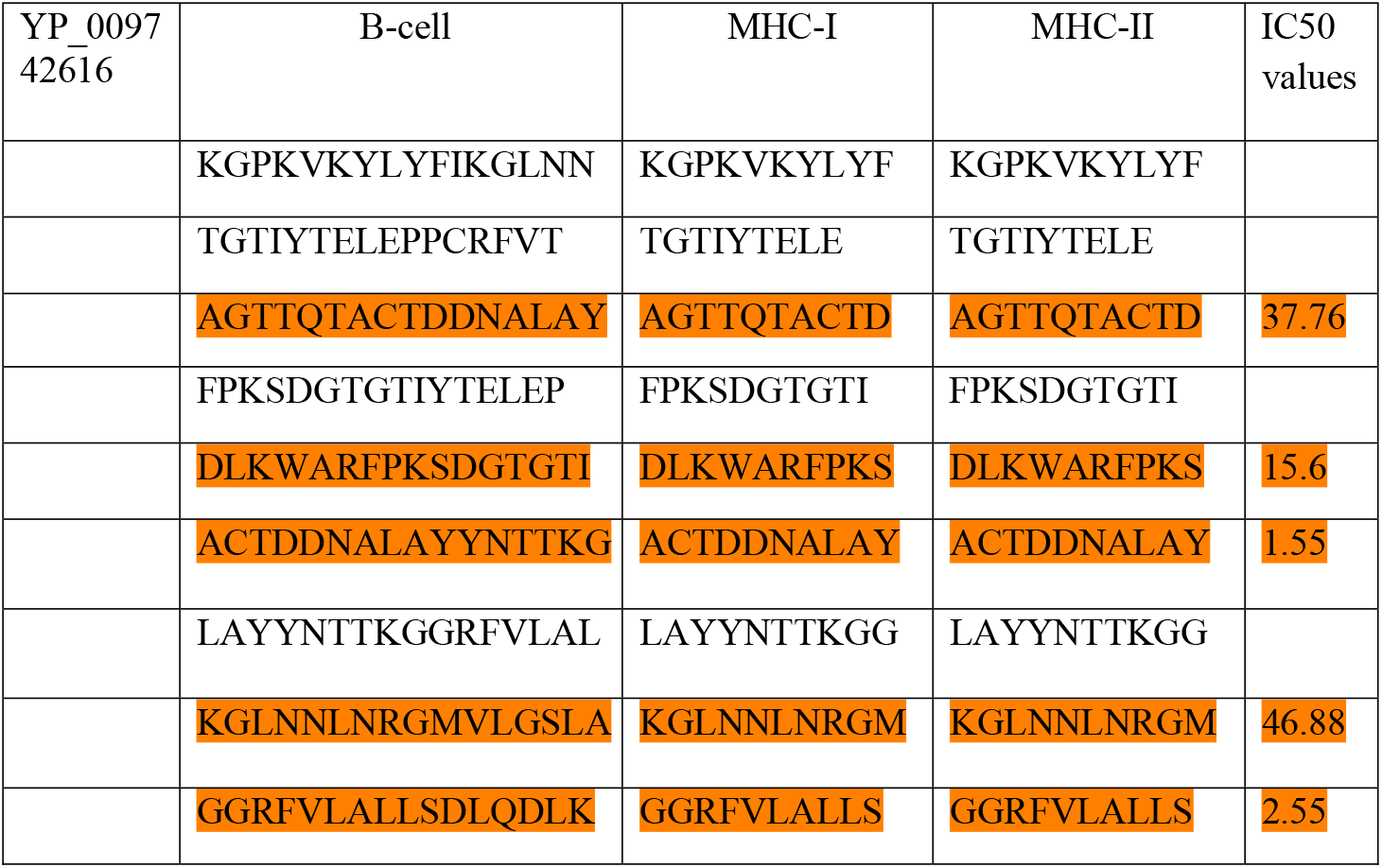
IC_50_ values of common epitopes.

### 3.3 Antigenecity

For the candidate protein sequences, antigenicity was evaluated using the Vaxijen server. Antigenecity is the ability of a chemical structure to bind explicitly with a group of products that have adaptive immunity. The VaxiJen 2.0 server uses an incipient alignment independent method for antigen prognostication predication. Protein sequences that showed antigenicity value (>0.4) were identified as antigenic and were subjected to further studies. Similarly, 11 proteins were predicted to be antigenic. The nsp9 protein was most antigenic (Table 1 & Supplementary Table1).

### 3.4 Human genome comparison

It is necessary to predict the similarity with human genome because as any similarity between human and virus genome may lead to side effects including auto immunity issues. Hence all 38 protein sequences were subjected to comparison with human proteome by using BLASTp tool with default parameters. “**No significant similarity found**” hits were considered as non-homologous proteins. All 11 proteins were found to be non-homologous to human proteome (Table 1 & Supplementary Table1).

### 3.5 Membrane topology analysis

The presence of these protein sequences either on the Transmembrane helix or outside the membrane were identified using the TMHMM server 2.0. This is necessary because proteins present in the core are less exposed and not viable. Hence the proteins which are present outside or in the Trans-membrane region were considered. In Table 1 & Supplementary Table1, all 11 proteins were shown to be present outside of trans membrane region.

### 3.6 Essentiality of protein sequences

The protein sequences were screened for essentiality using the DEG version 15.2 database which is a database that consists of the details of essential genes. Here for the submitted protein sequences BLAST is performed against the DEG database and hits are obtained which concludes that all 11 structural proteins are essential for the virus growth and survival. The protein sequences which fulfilled all the above mentioned filtration process were tabulated separately.

### 3.7. B-cell Epitope prediction

After obtaining final set of 11 proteins, B-cell epitopes were predicted for each of these sequences. B-cell epitope is a part of the protein that will bind with the antibody. Hence it is necessary to find the B-cell epitopes for these 11 proteins. The B-cell epitopes predicted for each 11 proteins are given in Supplementary Table 2.

### 3.8. T-cell Epitope prediction

Next we predicted MHC Class I and MHC Class II epitopes. All the epitopes are given in Supplementary Table 2. We looked for those epitopes that was common to both B-cell and Tcell epitopes. The list of common epitopes are given in Supplementary Table 2.

### 3.9. Inhibitory concentration

The common epitopes were checked for the inhibitory concentration (IC_50_) values that predicts epitope binding to major histocompatibility complexes (MHCs). For this, MHCpred tool was used. In this server common epitope was checked for its binding affinity with allele **DRB1*0101** as it is considered as immunodominant allele. All those common epitopes were selected whose IC_50_ value was <200 nM. Similarly, the epitopes with IC_50_ values less than 200 nM considered for epitope mapping on protein structure. We considered only those epitopes whose predicted IC_50_ values was less than 100 nM (Table2).

### 3.10. Prediction of 3-D structure and its validation

All proteins except spike protein (PDB ID: 6M0J and 6LVN) were modeled and validated (Supplementary Table 3)and used for epitope mapping. The final epitopes were found on six proteins namely Leader protein(nsp1)(Figure 1), 3c-like proteinase(nsp5)(Figure 2), Nsp8 (Figure 3), Nsp9 (Figure 4), Nsp9 (isoform)(Figure 5), Nsp10 (Figure 6) and Surface glycoprotein (spike protein).

After refinement the selected models were validated using SAVES server which helps in evaluating the dependability of the protein. SAVES version 5.0 server from NIH MBI laboratory is mostly useful for validation analysis. The quality was evaluated using VERIFY3D and PROCHECK (Supplementary Table 3). From this server, Ramachandran plot was also visualized for each protein (Figure 1-6). If a protein sequence is present in more disallowed region, then that protein is not considered.

### 3.11 Peptide mapping

Out of the 11 protein sequences, after validating the structure from the Ramachandran plot only 6 proteins (**nsp1, nsp5, nsp8, nsp9, nsp9(isoform) and nsp10**) were having final epitopes. The epitopes were mapped and again antigenicity of those epitopes were predicted. The eight surface exposed and high antigenecity epitopes (Fig 7 a–h) were selected for docking study with “DRB1*0101” (PDB ID “1AQD”).

**Fig 7a.**
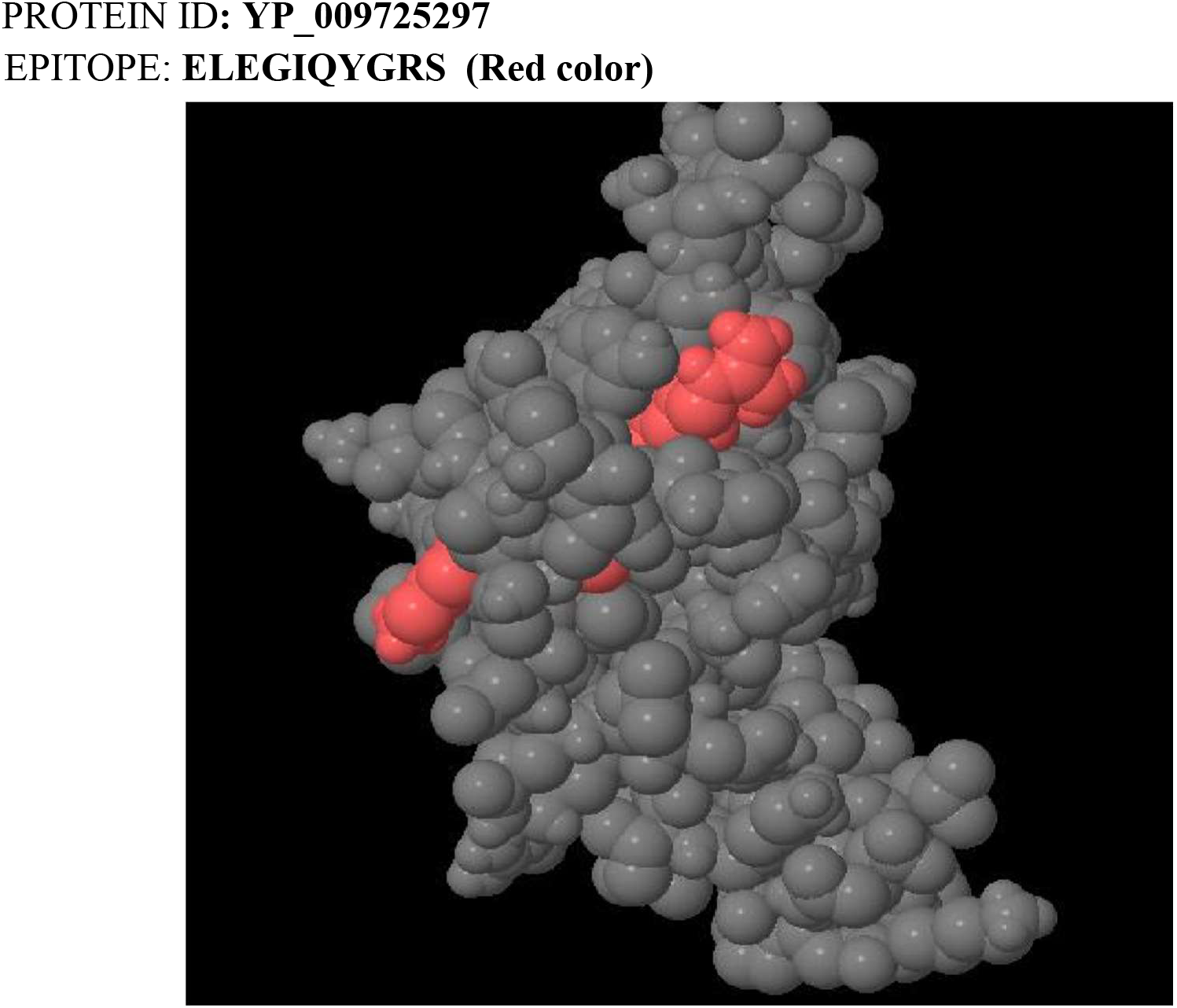
Position of epitope in YP_009725297.

**Fig 7b.**
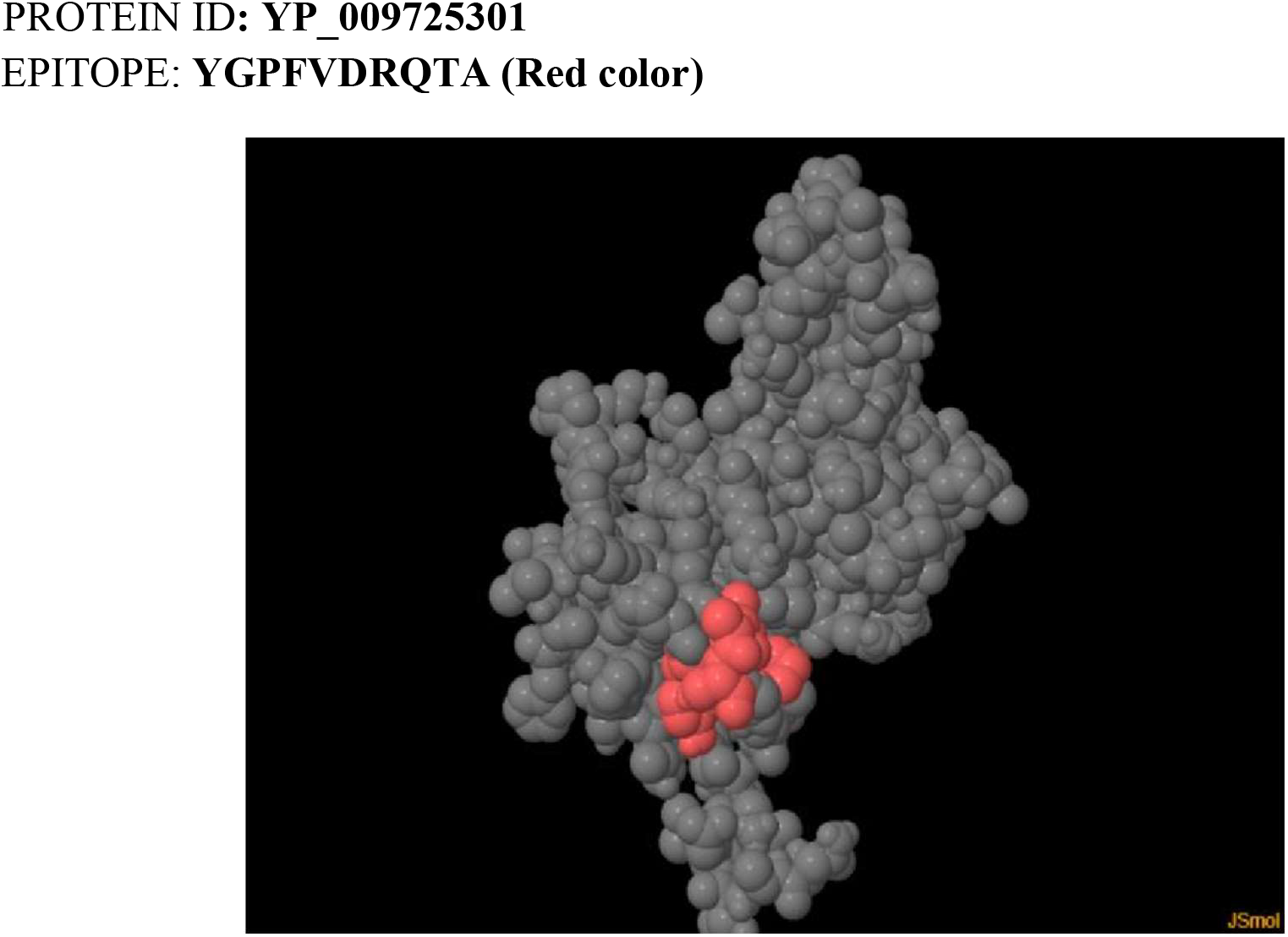
Position of epitope in YP_009725301.

**Fig 7c.**
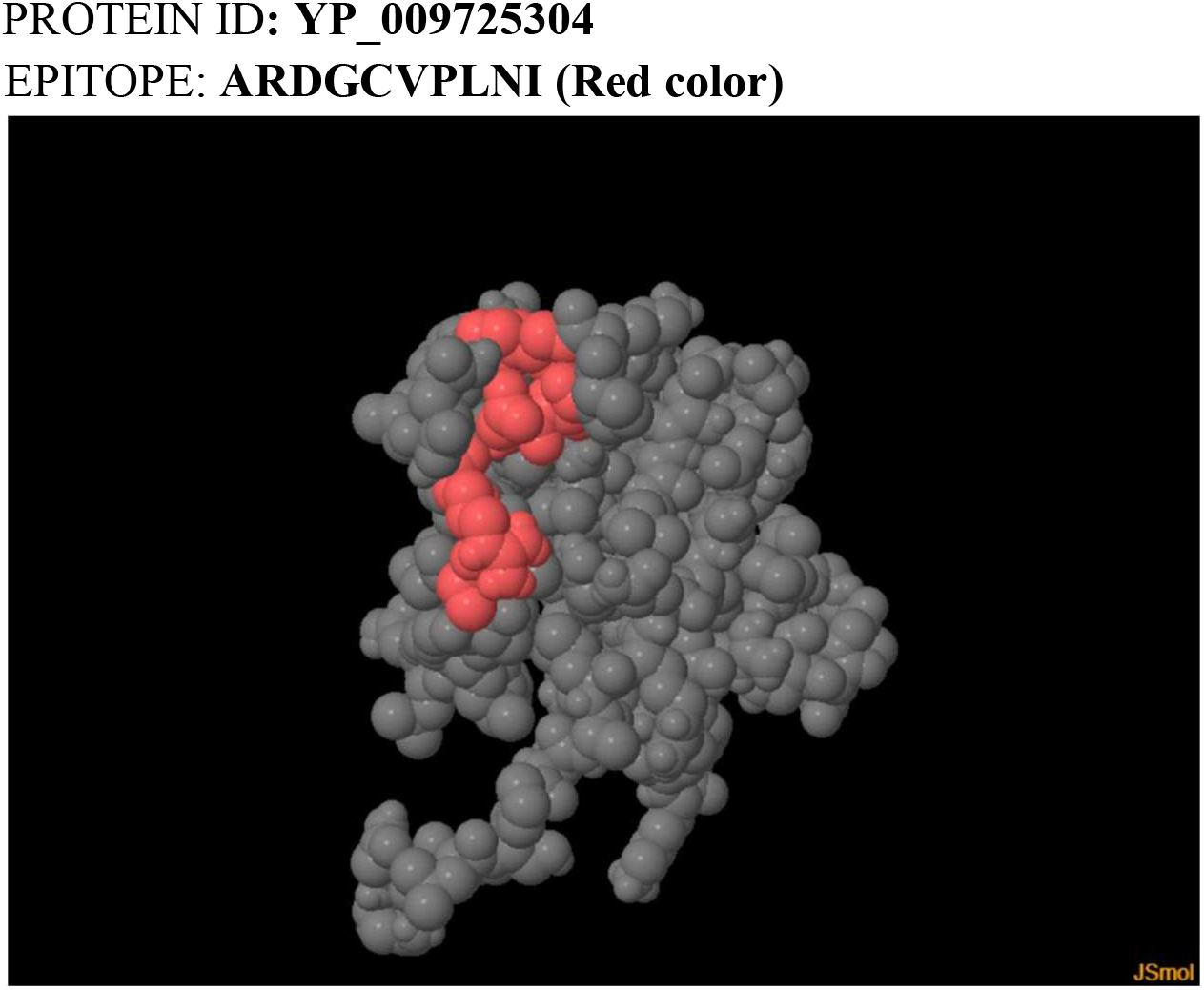
Position of epitope in YP_009725304.

**Fig 7d.**
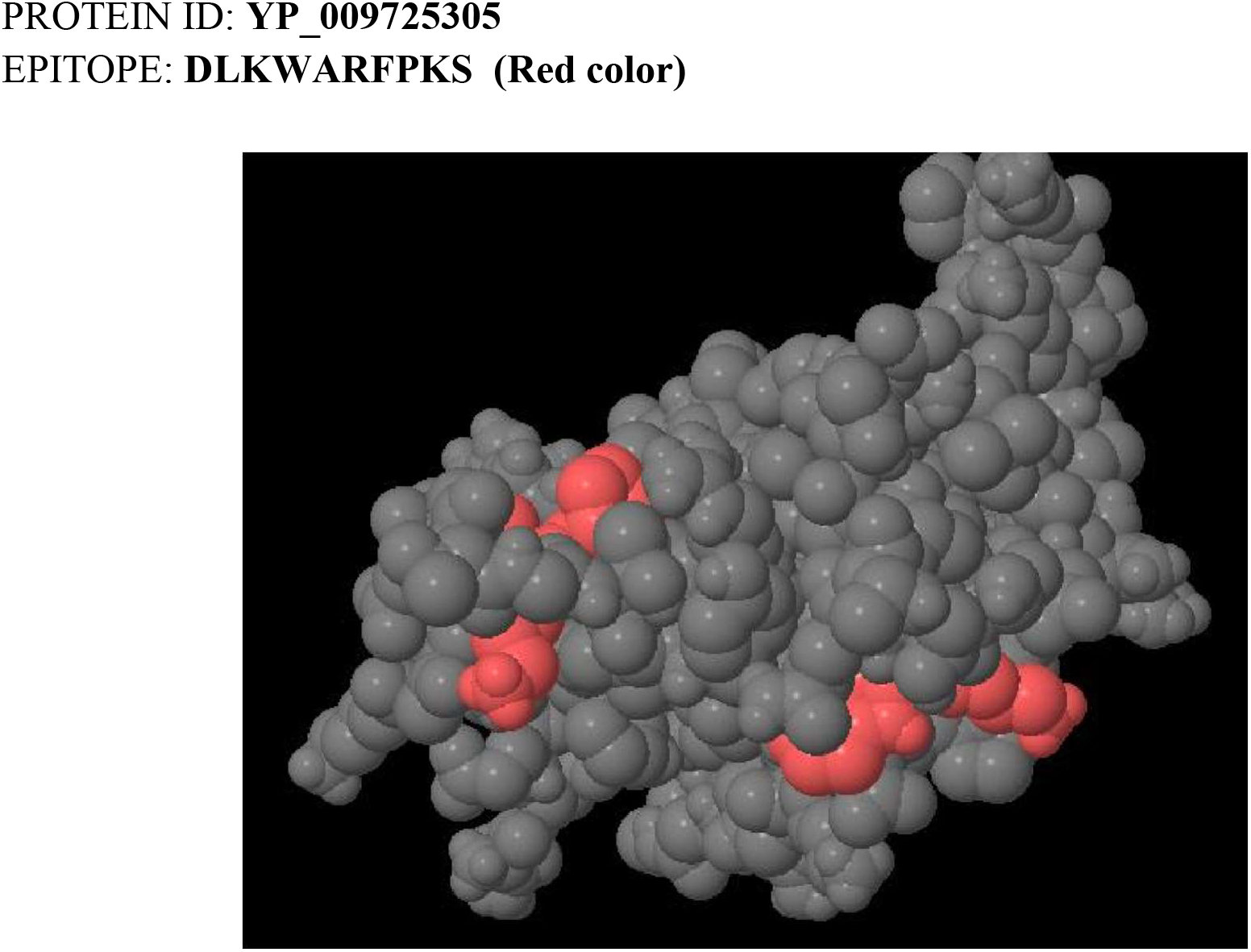
Position of epitope in YP_009725305.

**Fig 7e.**
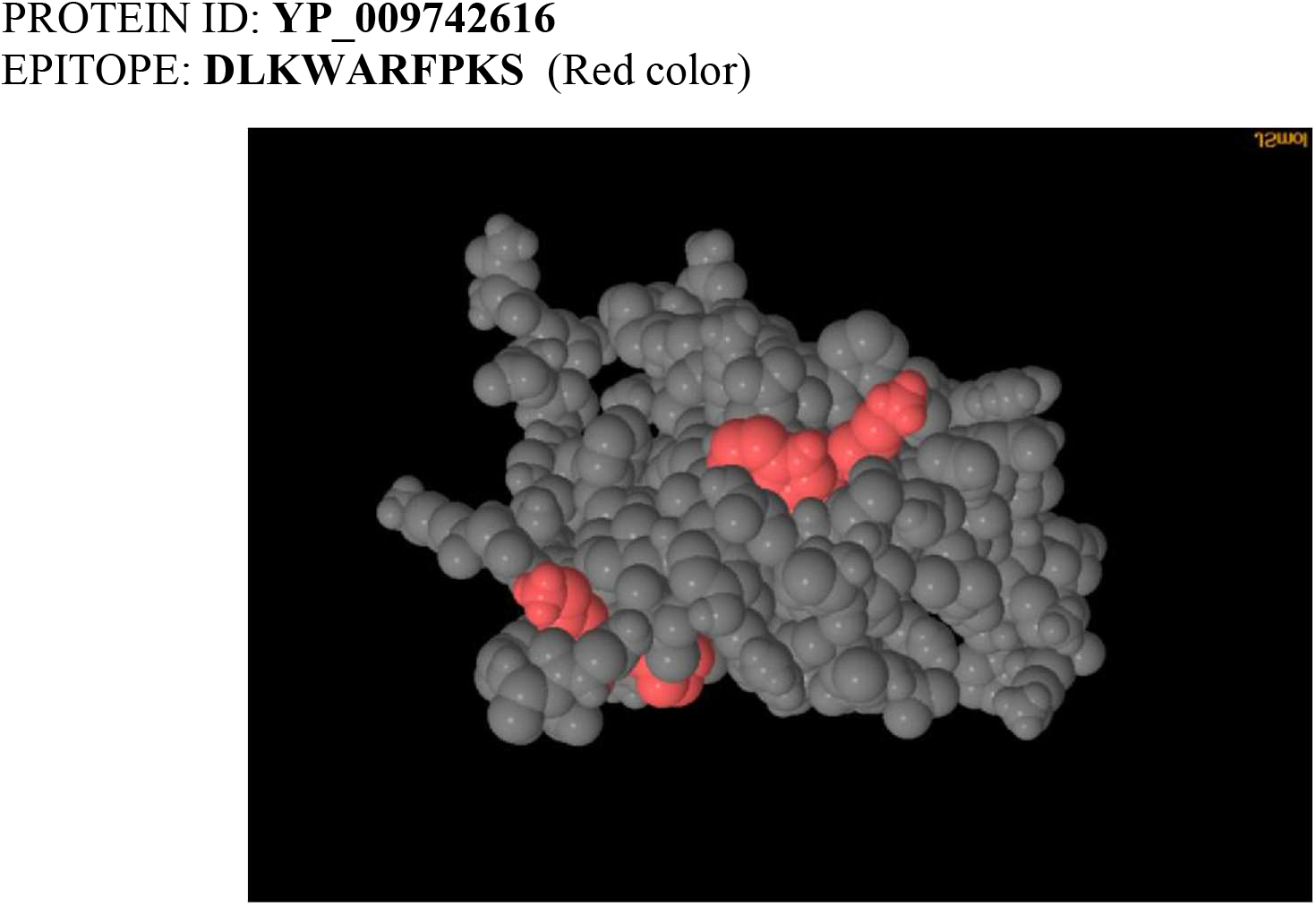
Position of epitope in YP_009742616.

**Fig 7f.**
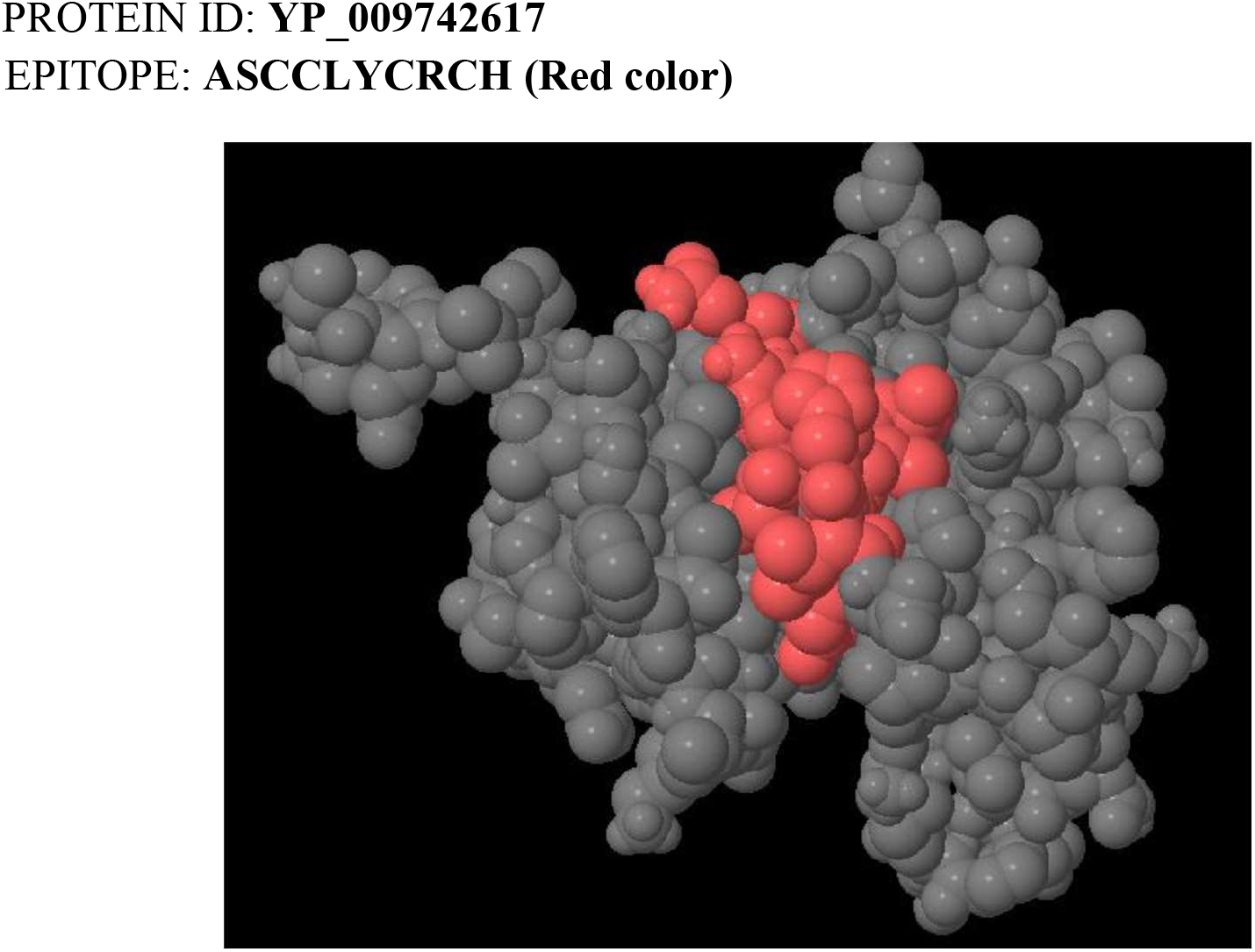
Position of epitope in YP_009742617.

**Fig 7g.**
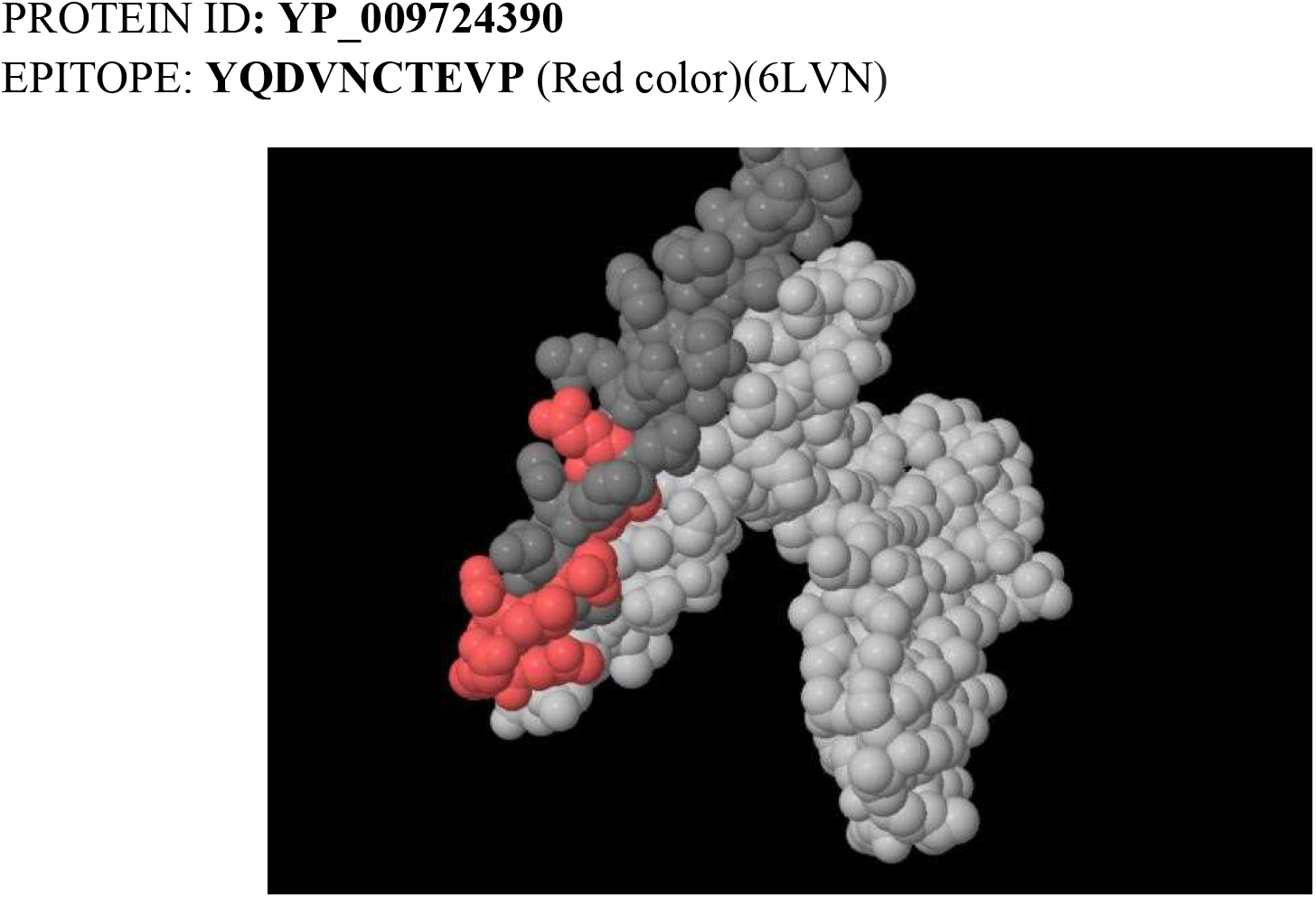
Position of epitope in YP_009724390.

**Fig 7h.**
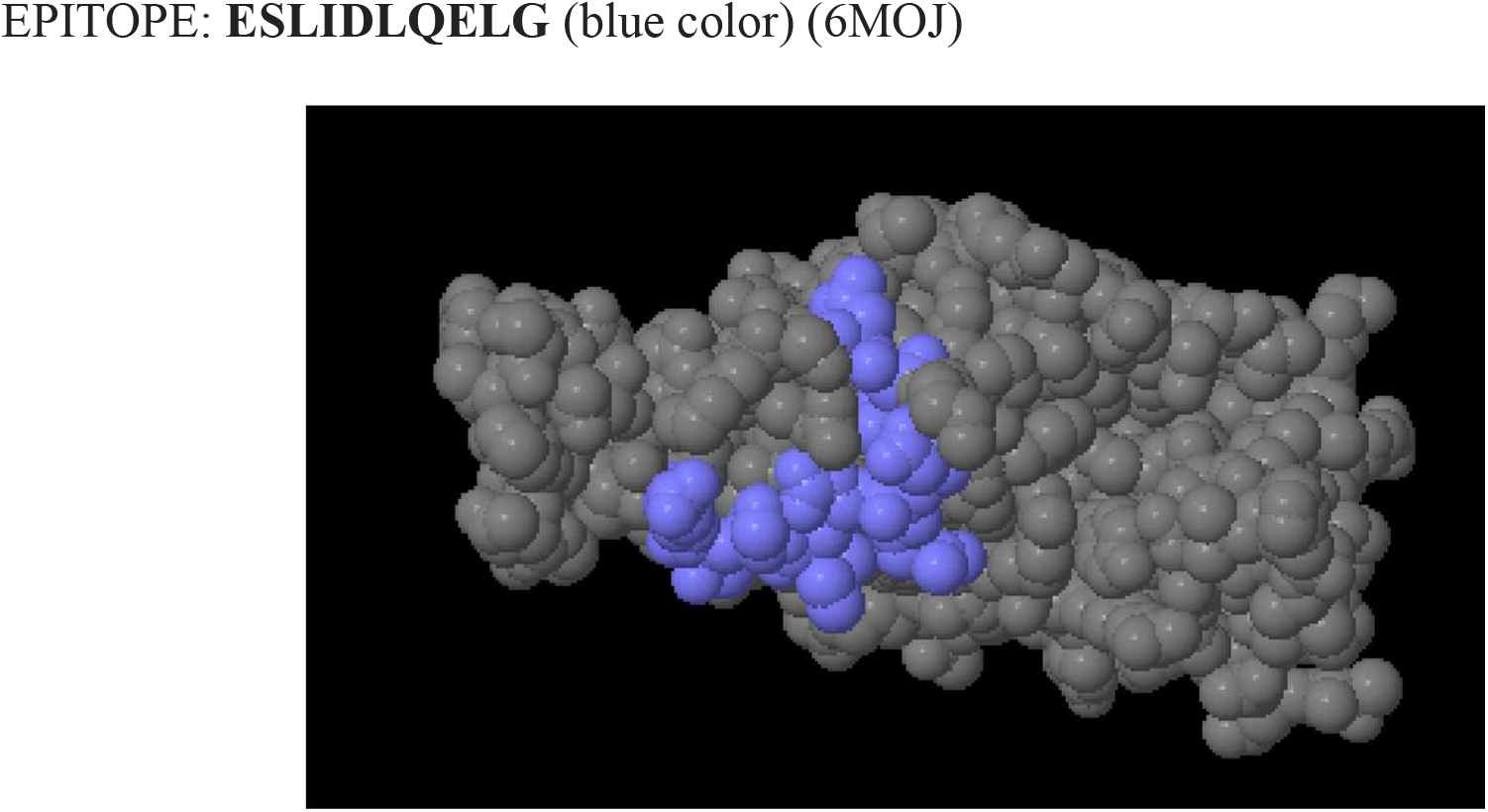
Position of epitope in YP_009724390.

### 3.12 Analysis of epitope binding and interaction

The best peptide-protein complexes were ranked based on interaction parameters which include hydrogen-bonding pattern, similarity score and accuracy score. For interpreting the binding mode of the complex Ligplot server was used and results are shown in Supplementary Figure file 1(Fig 8 a-g).

### 3.13 Analysis of hydrogen bonding pattern

From the obtained hydrogen-bonding interaction the one with the maximum number of hydrogen bonds were taken into consideration as this results in stability and more energetically favourable interactions. The epitopes with more number of hydrogen bonds (**ELEGIQYGRS and YGPFVDRQTA**) can be considered as more promising epitopes (Table 3).

**Table 3:**
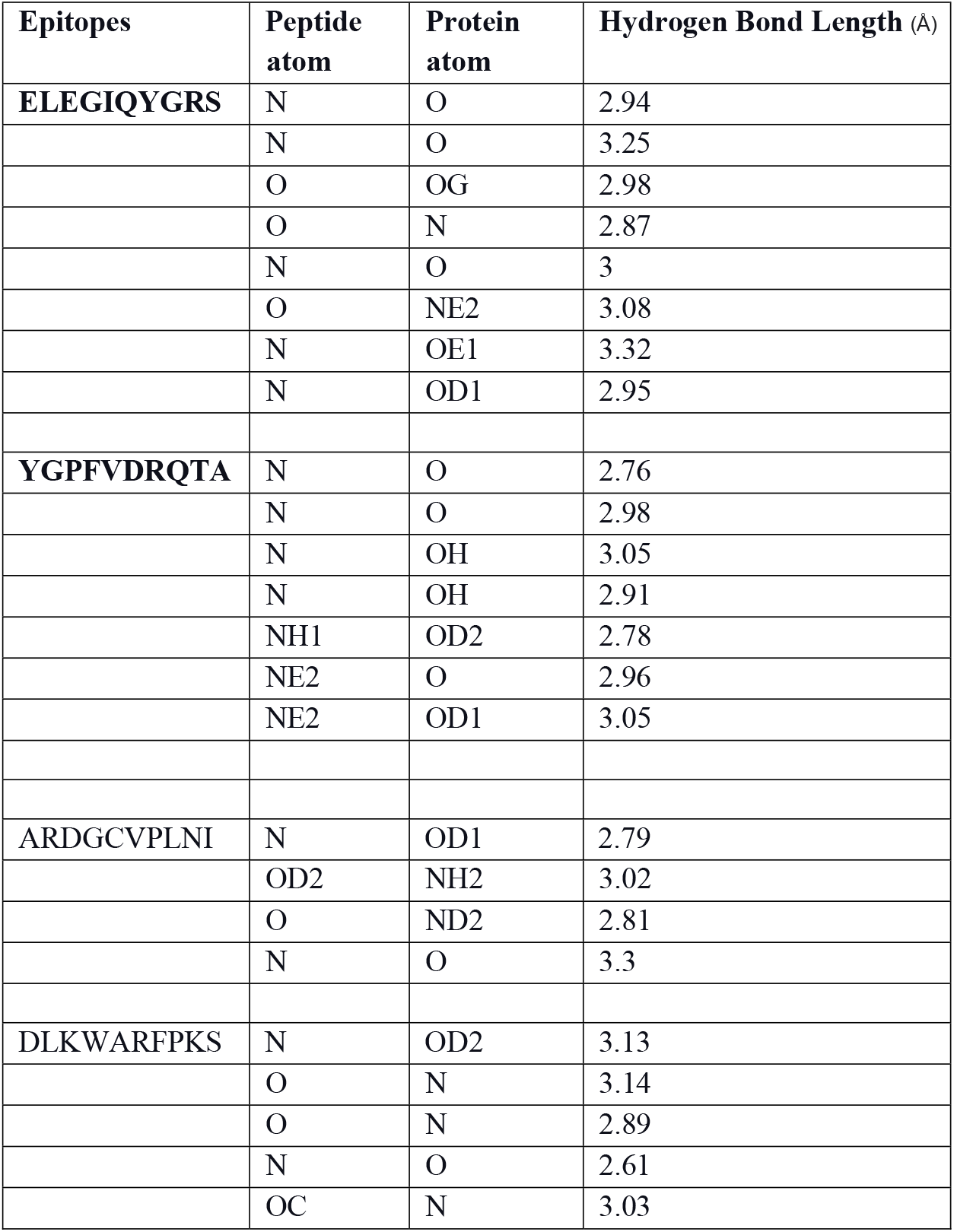
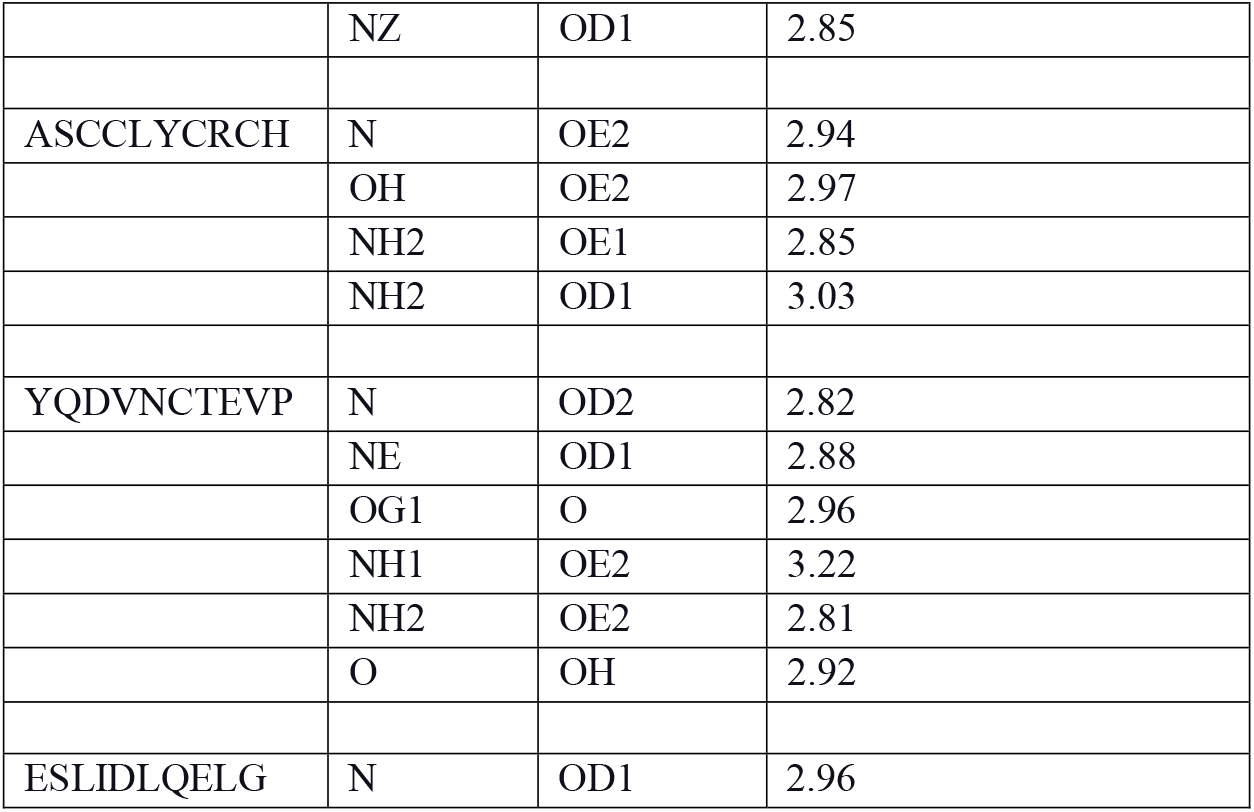
H bond between epitope atom and protein atom.

**ELEGIQYGRS** from Leader protein (NSP1), **YGPFVDRQTA** from 3c-like proteinase (nsp5), **DLKWARFPKS** from NSP9 and **YQDVNCTEVP** from Surface glycoprotein (spike protein) are the epitopes which has more hydrogen bonds. Hence these four epitopes could be considered as a more promising epitopes and these epitopes could be used for further clinical studies.

## CONCLUSION

Initially from the 38 protein sequences considered from the whole genome of SARS-CoV19, after fulfilling the necessary constraints which include antigenicity, virulency, nonhomologous to human proteome, presence of the epitope on the membrane and essentiality of protein only 11 protein sequences remained. For these 11 proteins linear B-cell and T-cell epitope prediction were done and the common epitopes were found. The protein structure for these sequences were modelled using I-TASSER and these models were subjected to refinement and validation processes after which only 6 protein were remaining. For the spike protein available protein model from PDB were taken into consideration. Now for the 7 protein models their epitope’s antigenicity were evaluated and the one with highest value for each protein was further considered for the docking process. These epitopes were docked with DRB1*0101 allele using GalaxyPepDock server and the docked structures obtained were analysed in LigPlot for hydrogen bonding interaction, similarity and accuracy score. As a result the hydrogen bonding pattern between the epitopes and the allele were analysed and the epitopes with more number of hydrogen bonds were considered to be more stable and energetically favourable. These epitopes could be used for further studies.

## Acknowledgement

We sincerely acknowledge SASTRA Deemed To be University for providing computational facility.

---

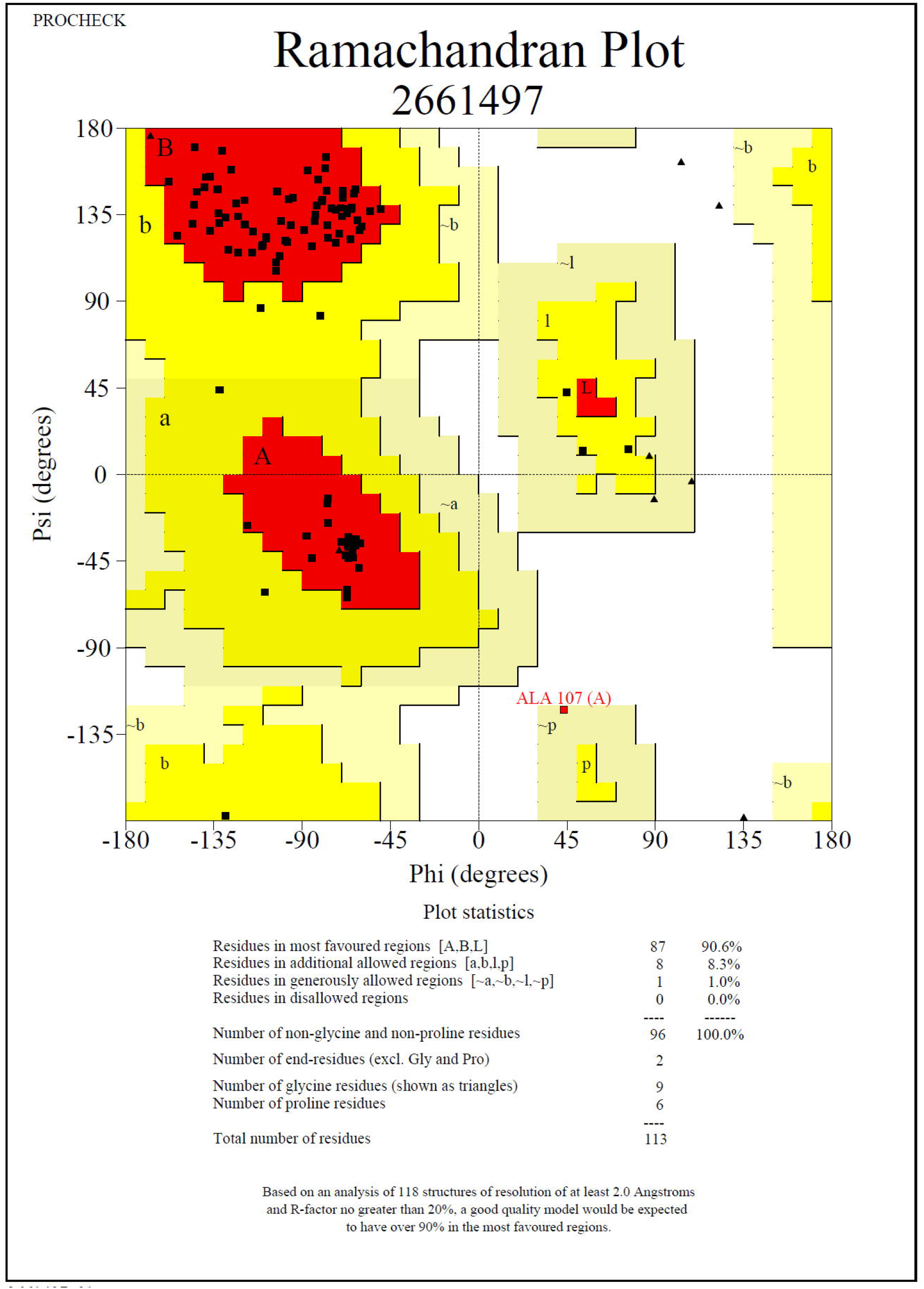

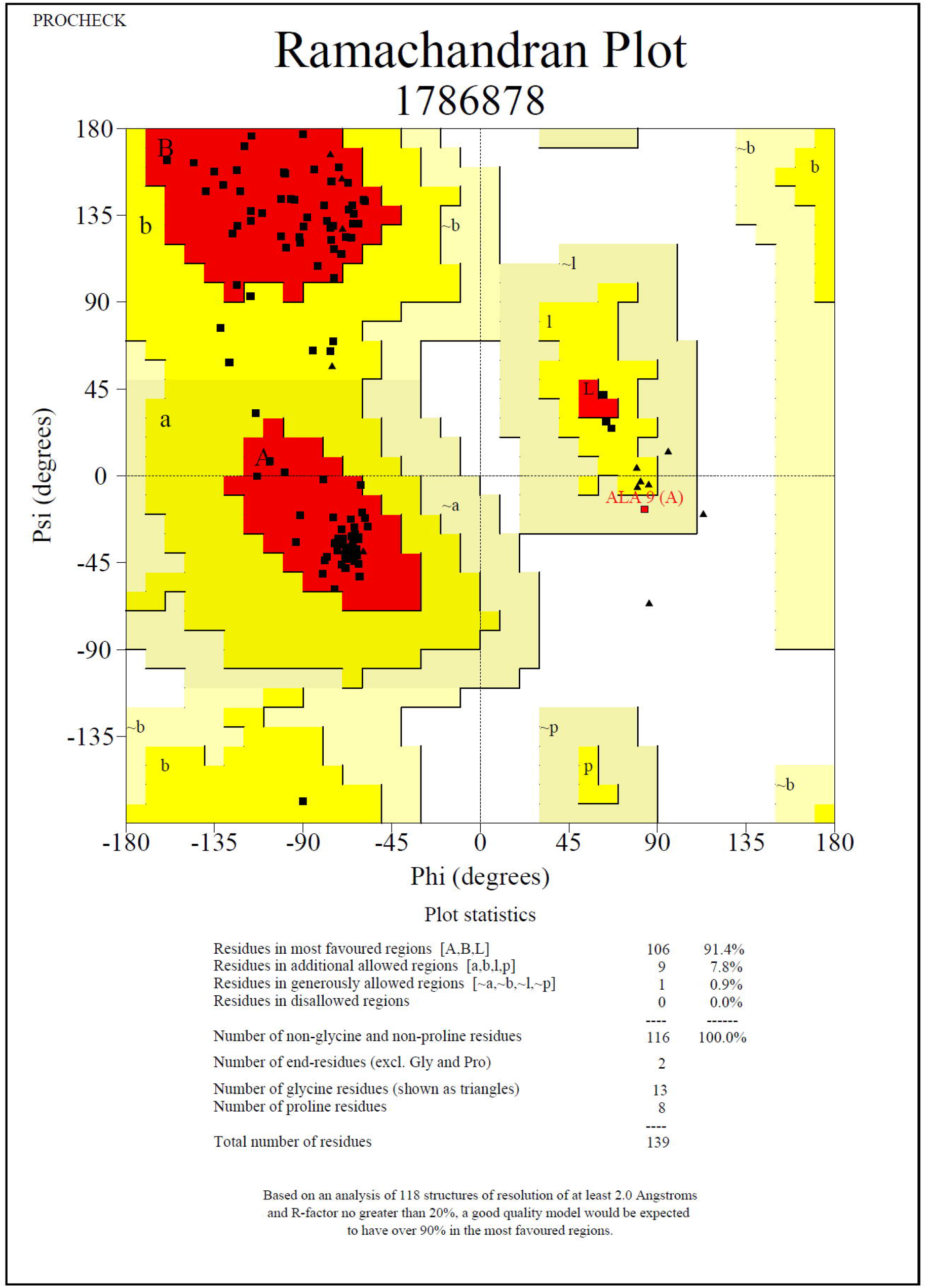

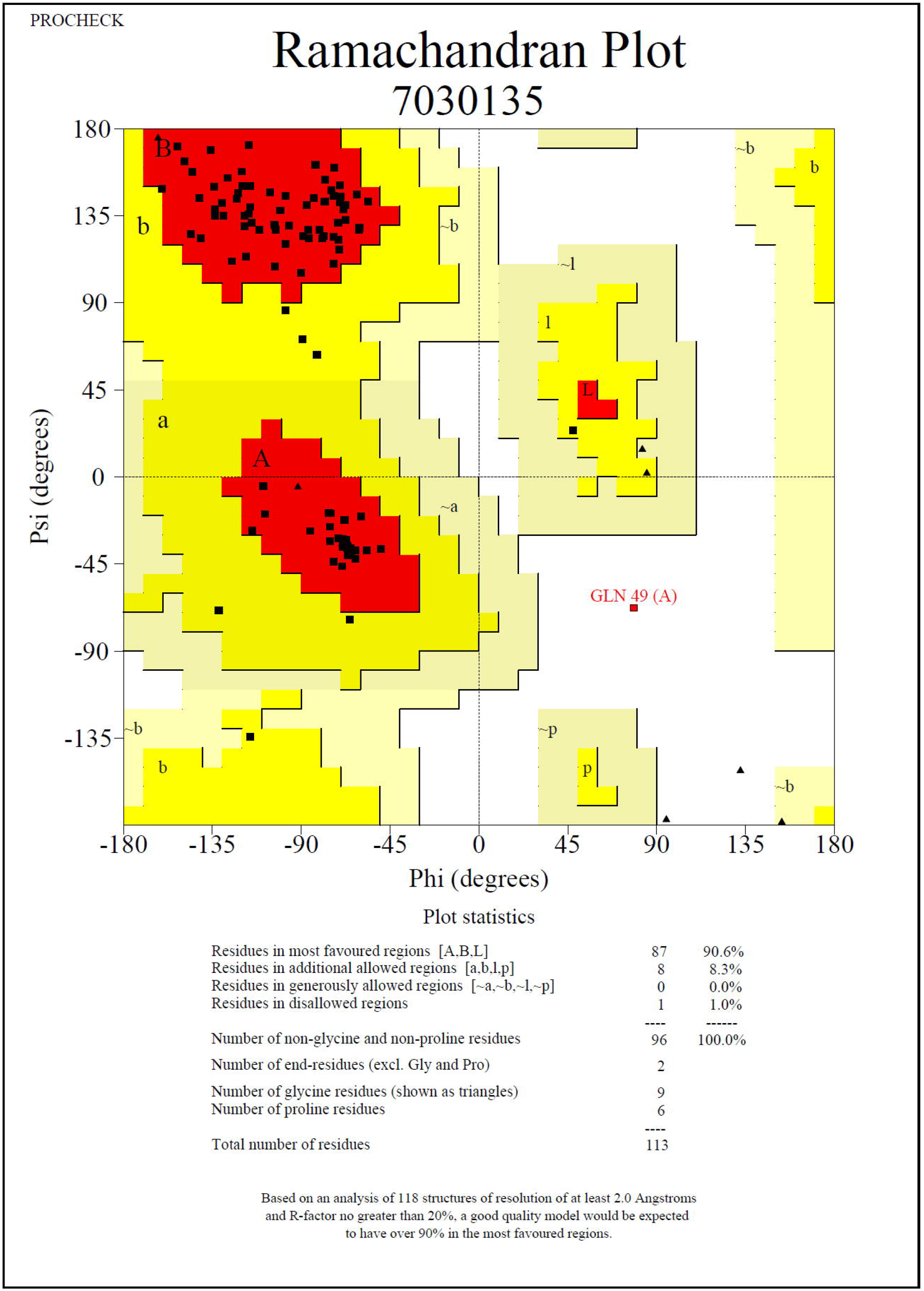

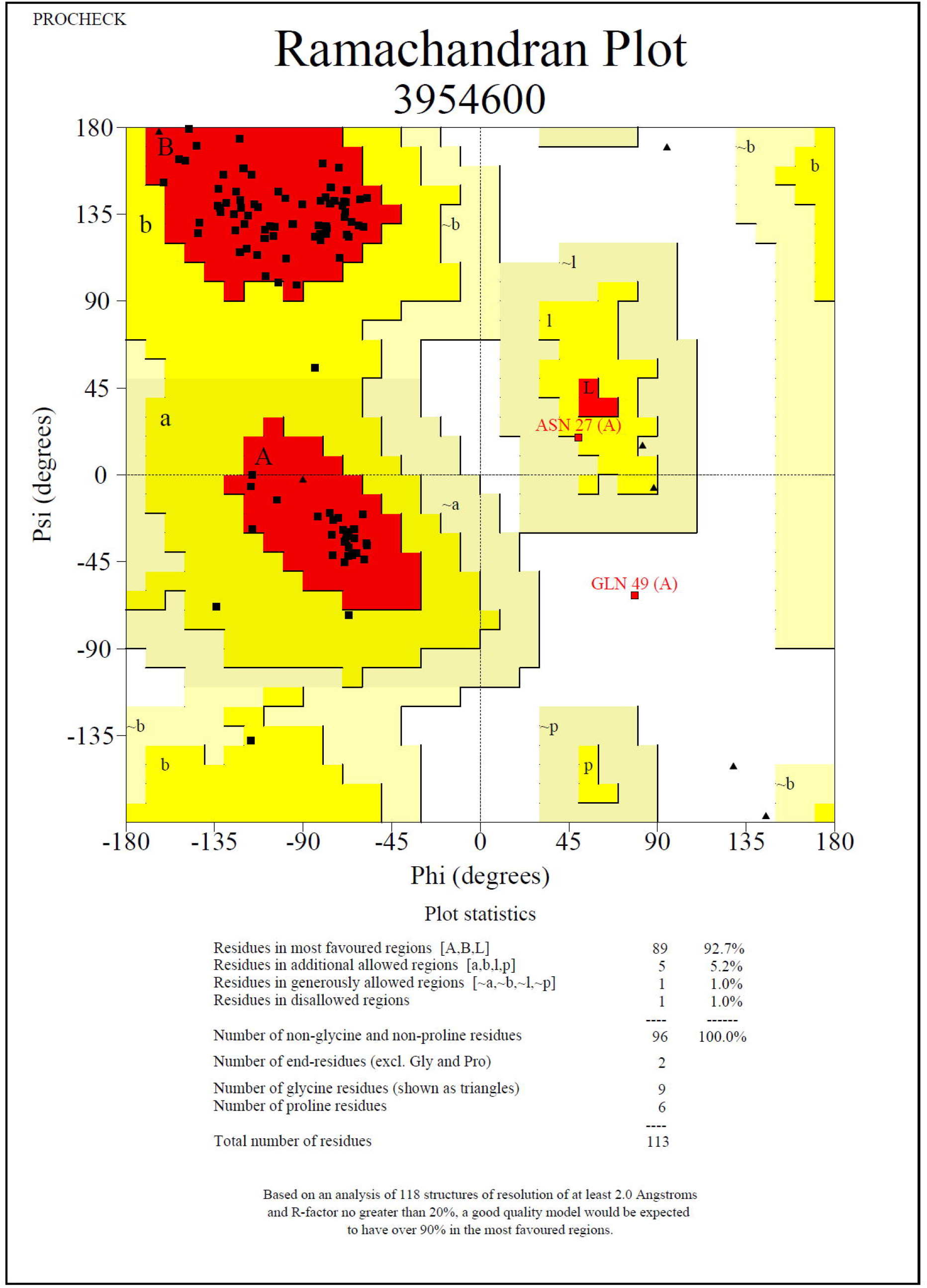

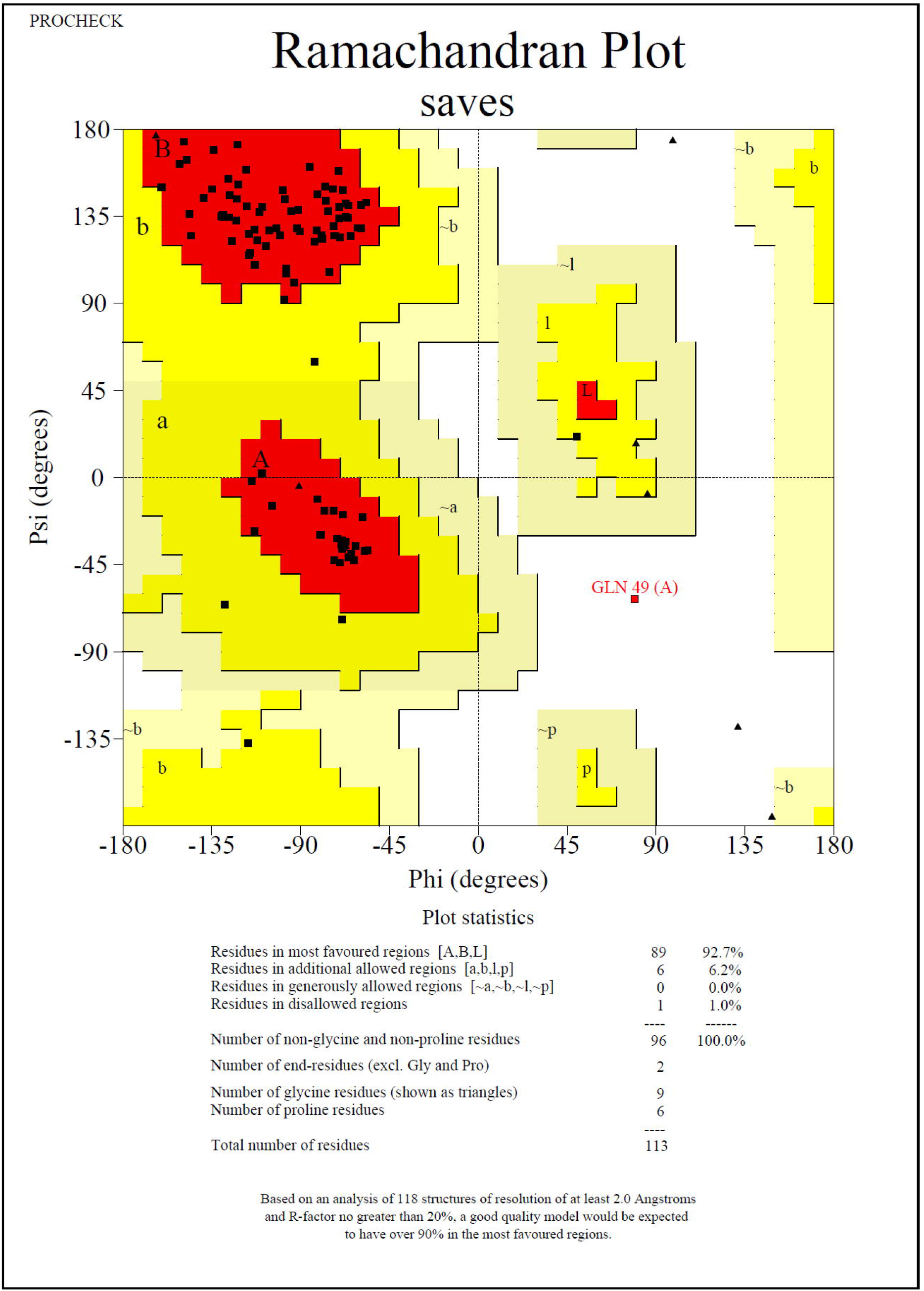

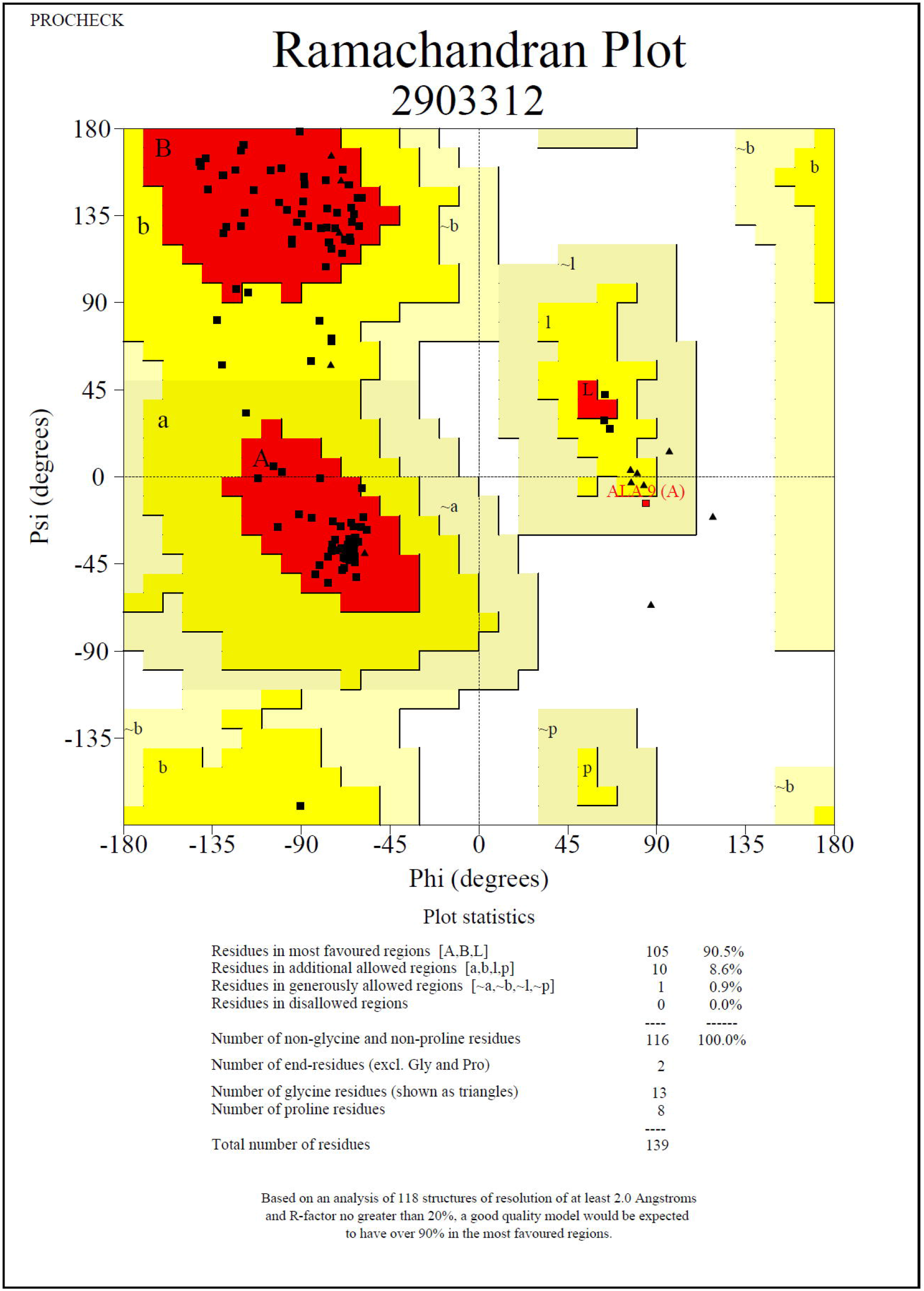

