## Supplementary figure 1 for "Peptide-based epitope design on non-structural proteins of SARS-CoV-2"

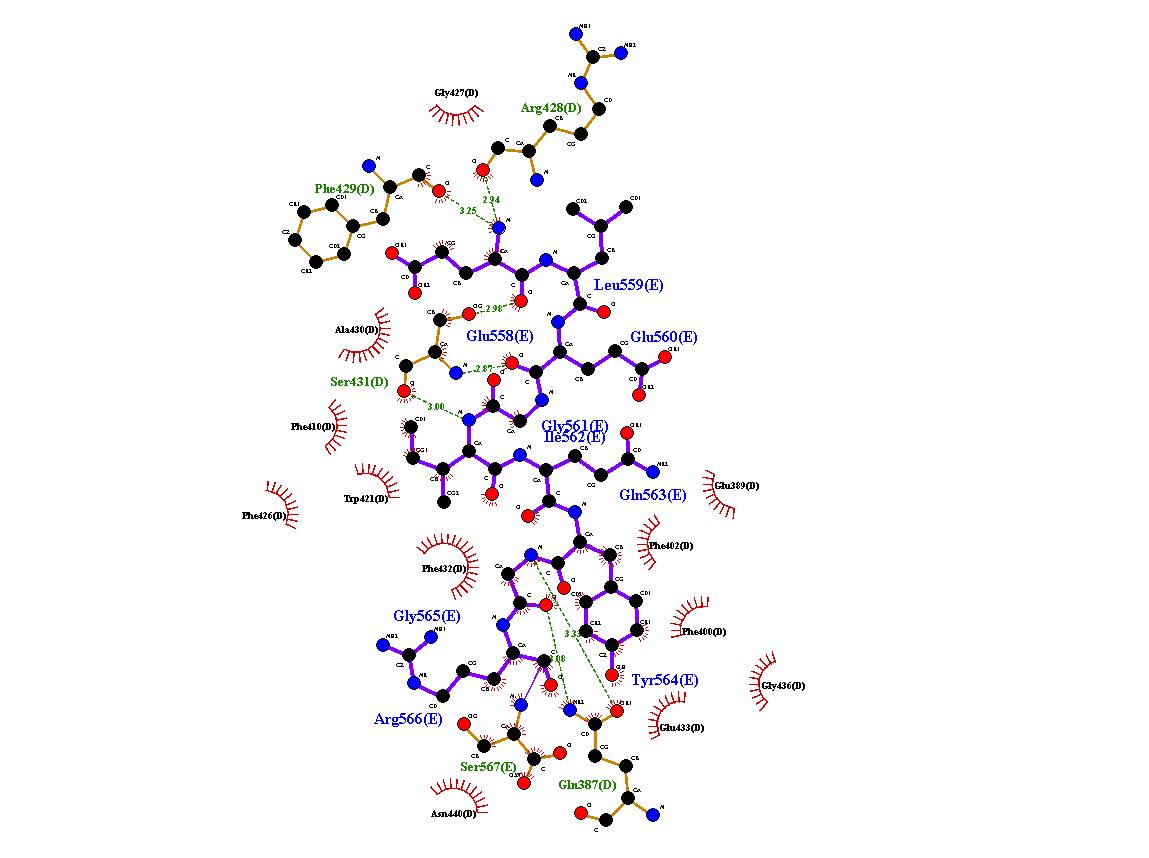
**Fig 8a Interaction of hydrogen bond between the protein (1AQD) and epitope ELEGIQYGRS (Blue color)**

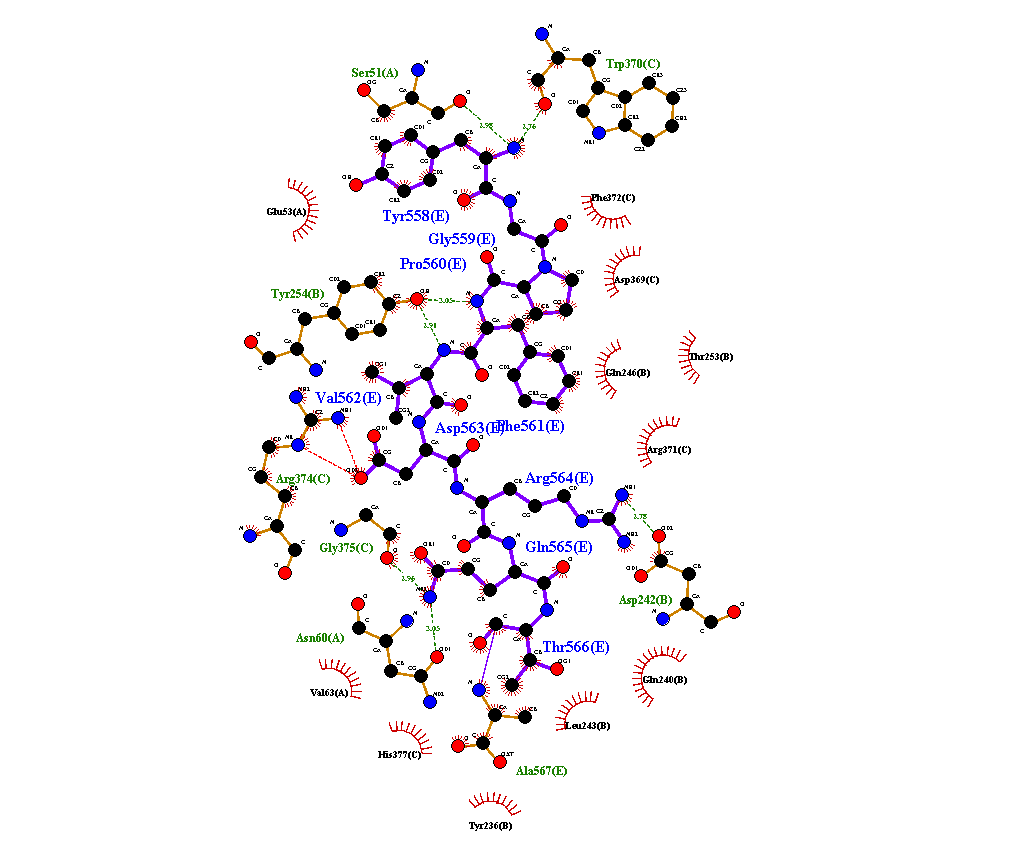

**Fig 8b Interaction of hydrogen bond between the protein (1AQD) and epitope YGPFVDRQTA**

**(Blue color)**

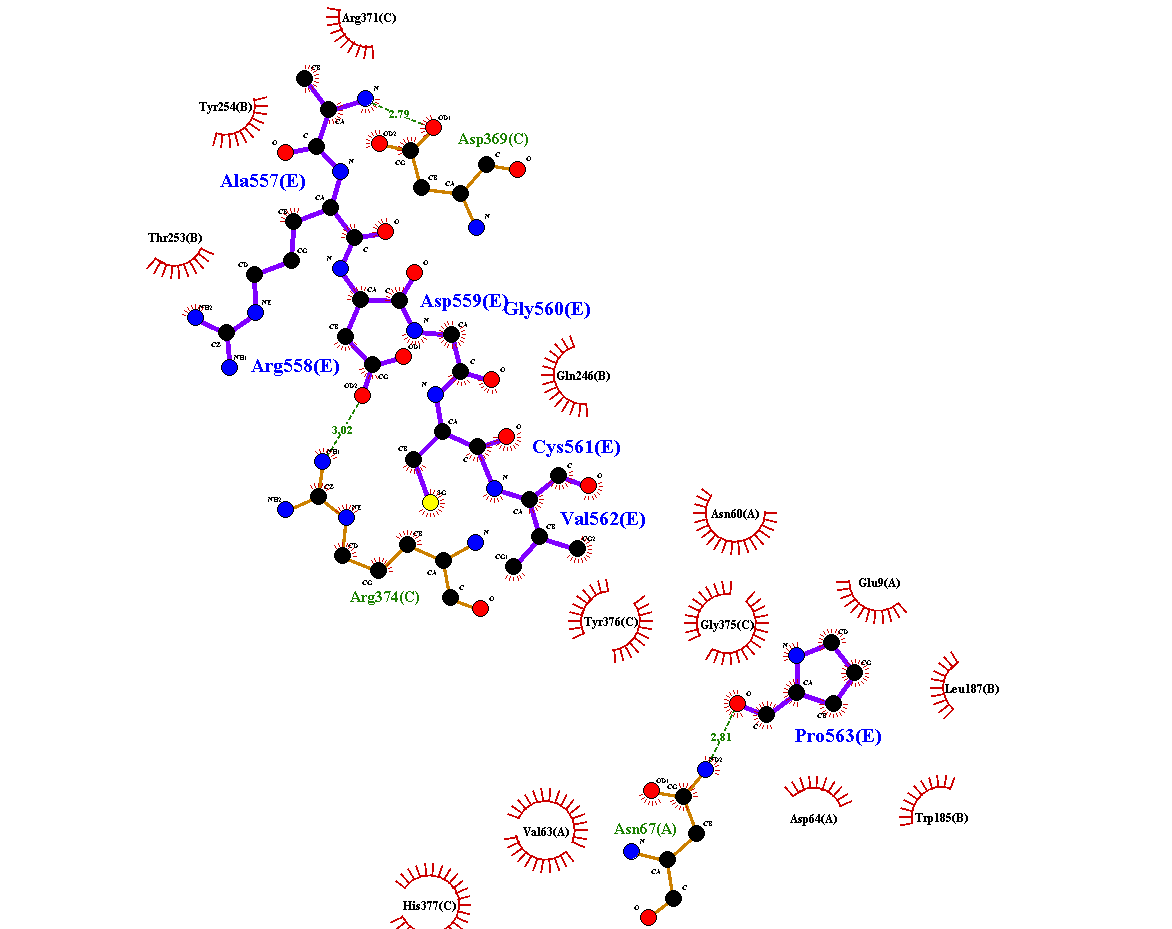
**Fig 8c Interaction of hydrogen bond between the protein (1AQD) and epitope ARDGCVPLNI**

**(Blue color)**

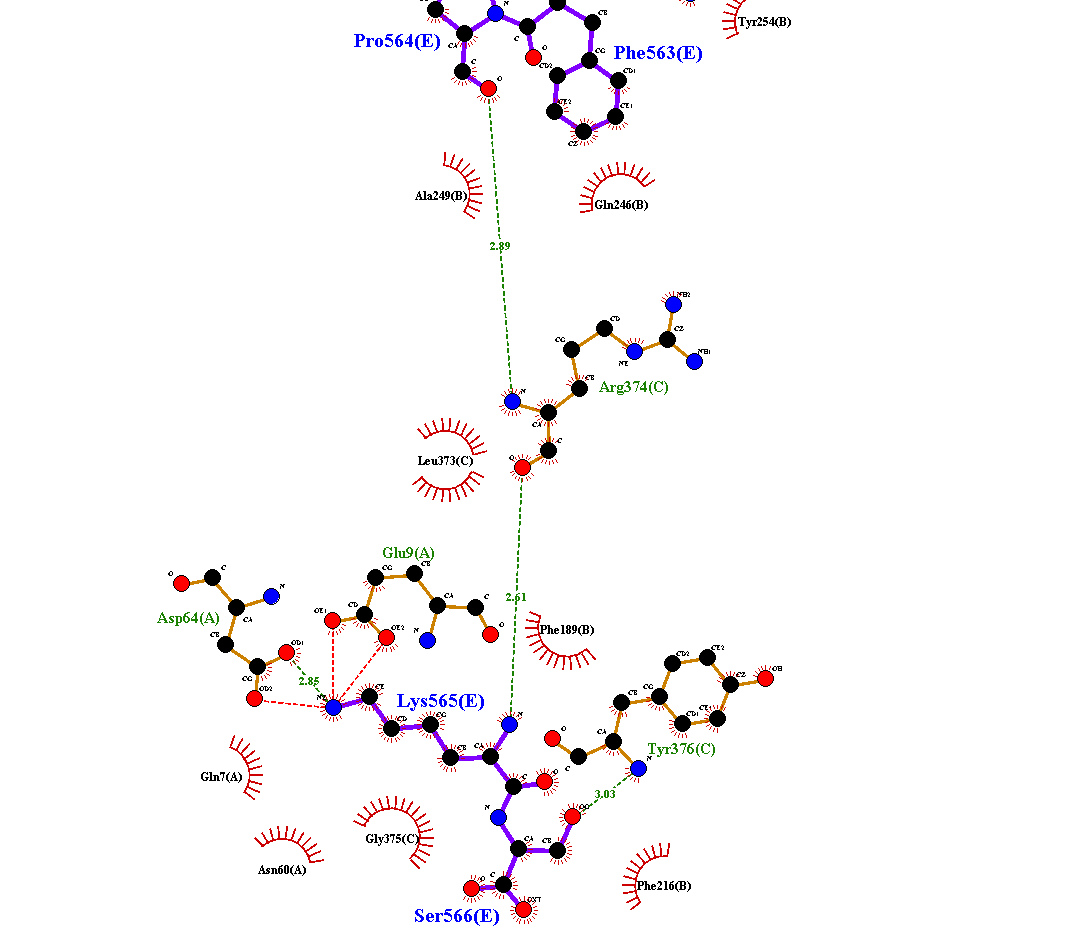

**Fig 8d Interaction of hydrogen bond between the protein (1AQD) and epitope DLKWARFPKS**

**(Blue color)**

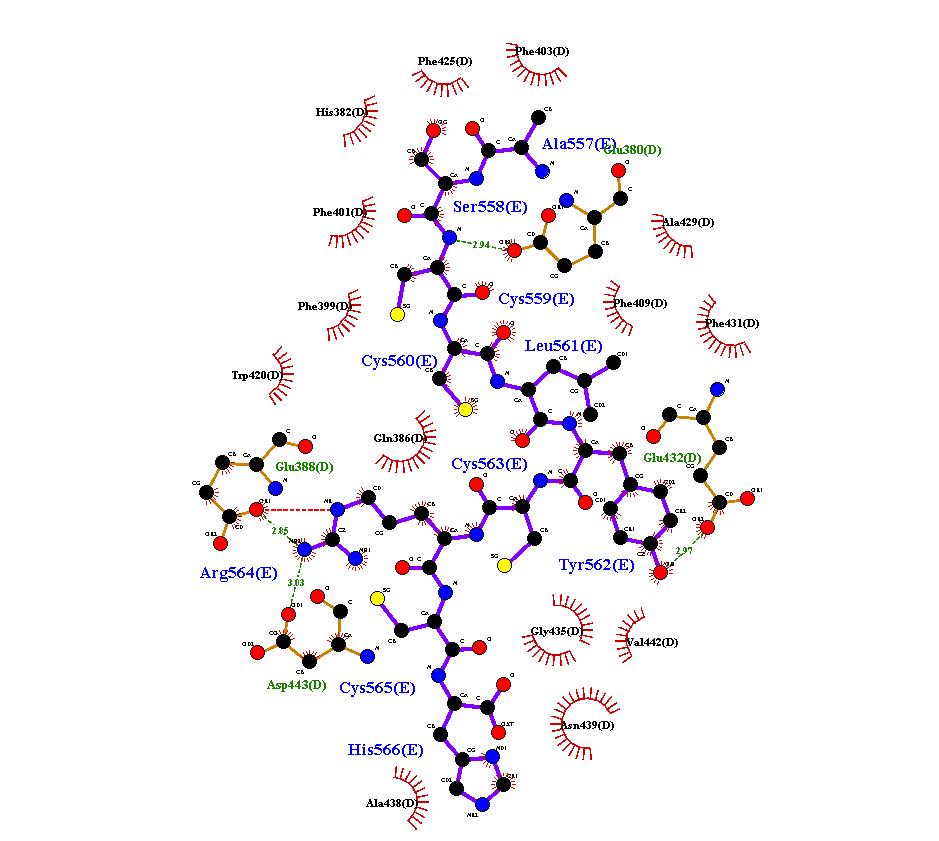
**Fig 8e Interaction of hydrogen bond between the protein (1AQD) and epitope ASCCLYCRCH**

**(Blue color)**

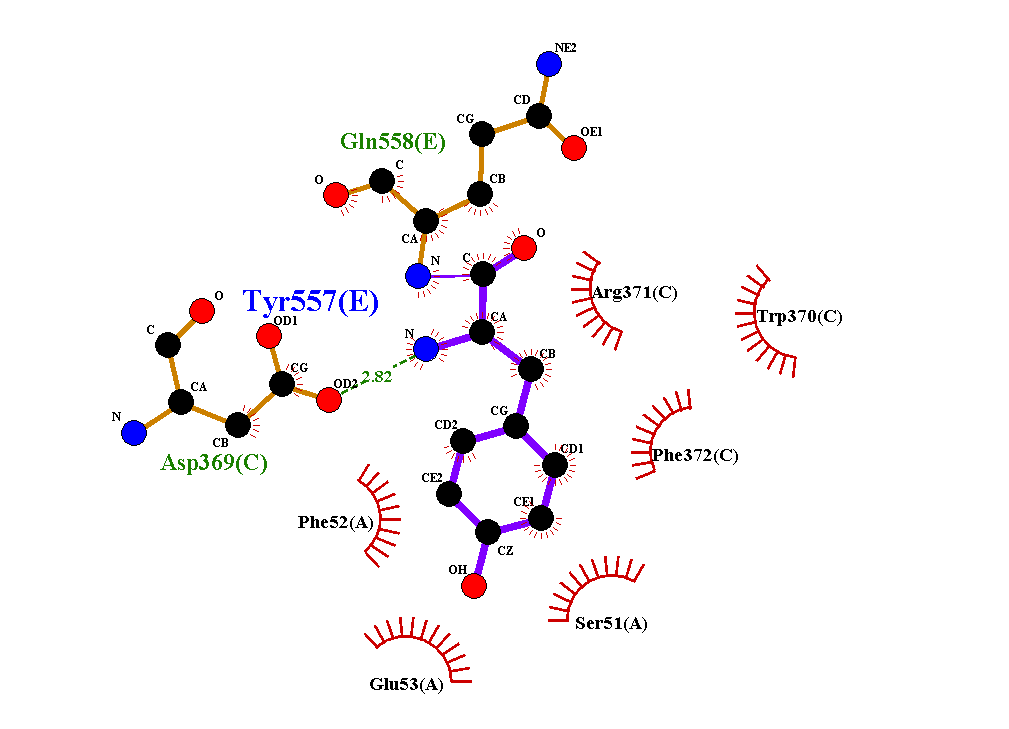

**Fig 8f Interaction of hydrogen bond between the protein (1AQD) and epitope YQDVNCTEVP**

**(Blue color)**

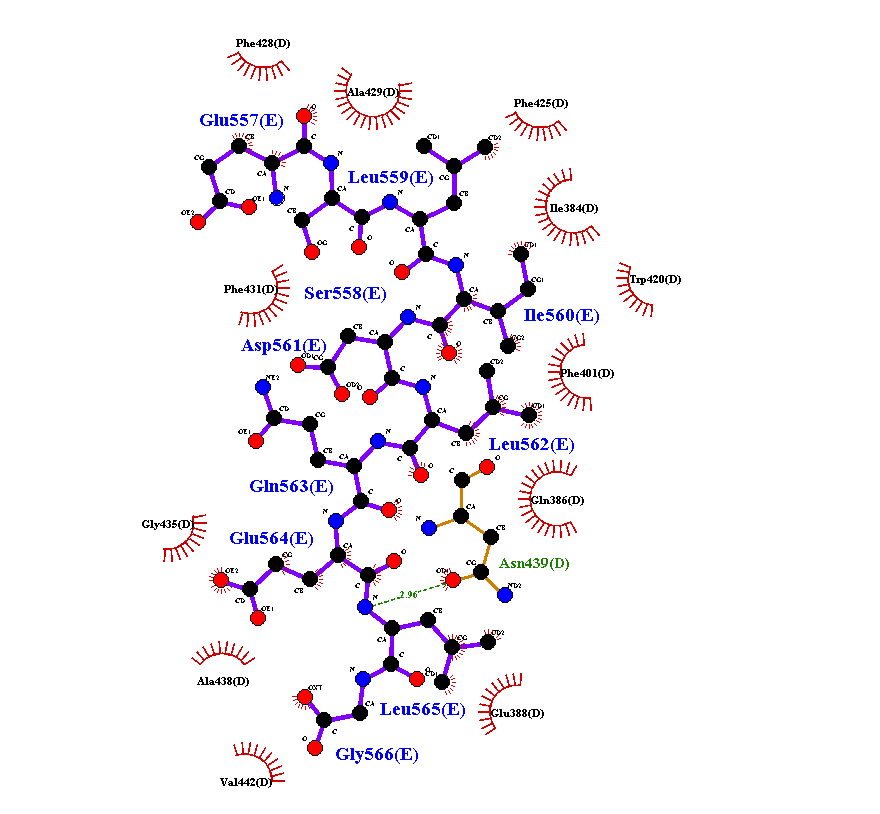
**Fig 8g Interaction of hydrogen bond between the protein (1AQD) and epitope ESLIDLQELG**

**(Blue color)**
