## Supplementary material for "Peptide-based epitope design on non-structural proteins of SARS-CoV-2": Table 1

**Table1: Final set of nCoV proteins for epitope prediction**

| **Locus** | **Protein name** | **Human**  **protein Similar**  **ity** | **Virule ncy** | **Antigeni**  **city** | **TM**  **Helix/Out side region** | **Essentia lity of protein** |
| --- | --- | --- | --- | --- | --- | --- |
| **YP_00972 5297** | Leader protein(nsp1) | No | P | 0.4064 | Outside | Yes |
| **YP_00972 5301** | 3c-like proteinase(ns p5) | No | P | 0.4159 | Outside | Yes |
| **YP_00972 5304** | nsp8 | No | P | 0.4008 | Outside | Yes |
| **YP_00972 5305** | nsp9 | No | P | 0.6476 | Outside | Yes |
| **YP_00972 5306** | nsp10 | No | P | 0.4039 | Outside | Yes |
| **YP_00974**  **2608** | Leader protein | No | P | 0.4064 | Outside | Yes |
| **YP_00974**  **2612** | 3c-like proteinase | No | P | 0.4078 | Outside | Yes |
| **YP_00974 2615** | nsp8 | No | P | 0.4107 | Outside | Yes |
| **YP_00974 2616** | nsp9 | No | P | 0.6226 | Outside | Yes |
| **YP_00974**  **2617** | nsp10 | No | P | 0.4149 | Outside | Yes |
| **YP_00972 4390** | Surface glycoprotein( spike protein) | No | P | 0.4661 | Outside/T M helix | Yes |
