## Supplementary material for "Peptide-based epitope design on non-structural proteins of SARS-CoV-2": Table 2

**Table 2: IC_50_ values of common epitopes**

| YP_0097  42616 | B-cell | MHC-I | MHC-II | IC_50_ values |
| --- | --- | --- | --- | --- |
|  | KGPKVKYLYFIKGLNN | KGPKVKYLYF | KGPKVKYLYF |  |
|  | TGTIYTELEPPCRFVT | TGTIYTELE | TGTIYTELE |  |
|  | AGTTQTACTDDNALAY | AGTTQTACTD | AGTTQTACTD | 37.76 |
|  | FPKSDGTGTIYTELEP | FPKSDGTGTI | FPKSDGTGTI |  |
|  | DLKWARFPKSDGTGTI | DLKWARFPKS | DLKWARFPKS | 15.6 |
|  | ACTDDNALAYYNTTKG | ACTDDNALAY | ACTDDNALAY | 1.55 |
|  | LAYYNTTKGGRFVLAL | LAYYNTTKGG | LAYYNTTKGG |  |
|  | KGLNNLNRGMVLGSLA | KGLNNLNRGM | KGLNNLNRGM | 46.88 |
|  | GGRFVLALLSDLQDLK | GGRFVLALLS | GGRFVLALLS | 2.55 |
