## Supplementary material for "Peptide-based epitope design on non-structural proteins of SARS-CoV-2": Table 3

**Table 3: H bond between epitope atom and protein atom**

| **Epitopes** | **Peptide atom** | **Protein atom** | **Hydrogen Bond Length** (Å) |
| --- | --- | --- | --- |
| **ELEGIQYGRS** | N | O | 2.94 |
|  | N | O | 3.25 |
|  | O | OG | 2.98 |
|  | O | N | 2.87 |
|  | N | O | 3 |
|  | O | NE2 | 3.08 |
|  | N | OE1 | 3.32 |
|  | N | OD1 | 2.95 |
| **YGPFVDRQTA** | N | O | 2.76 |
|  | N | O | 2.98 |
|  | N | OH | 3.05 |
|  | N | OH | 2.91 |
|  | NH1 | OD2 | 2.78 |
|  | NE2 | O | 2.96 |
|  | NE2 | OD1 | 3.05 |
| ARDGCVPLNI | N | OD1 | 2.79 |
|  | OD2 | NH2 | 3.02 |
|  | O | ND2 | 2.81 |
|  | N | O | 3.3 |
| DLKWARFPKS | N | OD2 | 3.13 |
|  | O | N | 3.14 |
|  | O | N | 2.89 |
|  | N | O | 2.61 |
|  | OC | N | 3.03 |
|  | NZ | OD1 | 2.85 |
| ASCCLYCRCH | N | OE2 | 2.94 |
|  | OH | OE2 | 2.97 |
|  | NH2 | OE1 | 2.85 |
|  | NH2 | OD1 | 3.03 |
| YQDVNCTEVP | N | OD2 | 2.82 |
|  | NE | OD1 | 2.88 |
|  | OG1 | O | 2.96 |
|  | NH1 | OE2 | 3.22 |
|  | NH2 | OE2 | 2.81 |
|  | O | OH | 2.92 |
| ESLIDLQELG | N | OD1 | 2.96 |
